# Heterozygous mutation to *Chd8* causes macrocephaly and widespread alteration of neurodevelopmental transcriptional networks in mouse

**DOI:** 10.1101/076158

**Authors:** Andrea L. Gompers, Linda Su-Feher, Jacob Ellegood, Tyler W. Stradleigh, Iva Zdilar, Nycole A. Copping, Michael C. Pride, Melanie D. Schaffler, M. Asrafuzzaman Riyadh, Gaurav Kaushik, Brandon Mannion, Ingrid Plajzer-Frick, Veena Afzal, Axel Visel, Len A. Pennacchio, Diane Dickel, Jason P. Lerch, Jacqueline N. Crawley, Konstantinos S. Zarbalis, Jill L. Silverman, Alex S. Nord

## Abstract

The chromatin remodeling gene *CHD8* represents a central node in early neurodevelopmental gene networks implicated in autism. We examined the impact of heterozygous germline *Chd8* mutation on neurodevelopment in mice. Network analysis of neurodevelopmental gene expression revealed subtle yet strongly significant widespread transcriptional changes in *Chd8*^+/−^ mice across autism-relevant networks from neurogenesis to synapse function. *Chd8*^+/−^ expression signatures included enrichment of RNA processing genes and a Chd8-regulated module featuring altered transcription of chromatin remodeling, splicing, and cell cycle genes. *Chd8*^+/−^ mice exhibited increased proliferation during brain development and neonatal increase in cortical length and volume. Structural MRI confirmed regional brain volume increase in adult *Chd8*^+/−^ mice, consistent with clinical macrocephaly. Adult *Chd8*^+/−^ mice displayed normal social interactions, and repetitive behaviors were not evident. Our results show that *Chd8*^+/−^ mice exhibit neurodevelopmental changes paralleling *CHD8*^+/−^ humans and show that *Chd8* is a global genomic regulator of pathways disrupted in neurodevelopmental disorders.

## Introduction

DNA packaging determines the transcriptional potential of a cell and is central to the development and function of metazoan cell types. Chromatin remodeling complexes control local chromatin state, yielding either transcriptional activation or repression. Pluripotency, proliferation, and differentiation are dependent on genomic regulation at the chromatin level, and proteins that control chromatin packaging are critical in development and cancer^1^. Although many chromatin remodeling factors function across systems, case sequencing efforts have linked mutations of chromatin genes with specific, causal roles in neurodevelopmental disorders^2–5^. This finding is particularly strong for rare and *de novo* mutations in autism spectrum disorder (ASD)^6,7^. Understanding how mutations to chromatin remodeling genes affect transcriptional regulation during brain development may reveal developmental and cellular mechanisms driving neurodevelopmental disorders.

A key gene that has emerged from studies profiling rare and *de novo* coding variation in ASD is the chromatin remodeler, *CHD8* (*Chromodomain helicase DNA binding protein 8*)^8^. In addition to ASD, *CHD8*^+/−^ individuals exhibit macrocephaly, distinct craniofacial morphology, mild-to-severe cognitive impairment, and gastrointestinal problems^8^. *CHD8* mutation also has been linked to attention deficit hyperactivity disorder, seizures, and schizophrenia^4,8^, as well as cancer^9,10^. Homozygous deletion of *Chd8* in mice is early embryonic lethal^11^. *Chd8* knockdown in zebrafish recapitulated macrocephaly and gastrointestinal phenotypes^8,12^, suggesting a high degree of evolutionary conservation of Chd8 function in brain development.

Studies of genetic and protein networks have raised the possibility that CHD8 is a central node and master regulator of early neurodevelopmental networks implicated in autism^12–14^. CHD8 has been proposed to achieve this regulatory function in brain development by binding to relevant gene promoters and enhancers^12,14^. CHD8 DNA binding and knockdown studies in human and mouse tissues and cells have revealed a multitude of genes directly and indirectly activated or repressed by CHD8 during neurodevelopment^12,14^.

Consequently, characterizing the functional impact of heterozygous *CHD8* mutation on brain development could reveal generalizable mechanisms linking chromatin biology to pathology. Towards this goal, we generated two new *Chd8* mutant mouse lines and performed analyses to characterize neuroanatomic, transcriptional, and behavioral phenotypes of *Chd8*^+/−^ mice. Our interrogations identified changes in structural and developmental neuroanatomy and subtle but highly significant changes to developmental gene expression. These results provide insight into *in vivo* pathological changes, showing that germline *Chd8* haploinsufficiency results in altered gene expression across neurodevelopment and produces increased regional brain volume. The results from this study indicate the presence of genomic and neuroanatomic phenotypes that parallel the clinical signature of human *CHD8* mutations, suggesting similar neurodevelopmental pathology between human and mouse.

## Results

### Mice harboring heterozygous germline Chd8 mutation exhibit megalencephaly

We used CRISPR/Cas9 targeting to generate two mouse lines harboring short deletions in *Chd8*, upstream of the majority of human mutations identified in autism cohorts^8^ (Figure 1A-1C). Consistent with an earlier study^11^, our two newly generated *Chd8* alleles (5 and 14 bp deletions within exon 5) resulted in embryonic lethality in homozygous mutants, but heterozygous (*Chd8*^+/−^) mice were viable, reached a normal lifespan, and were fertile irrespective of sex. Quantitative PCR (qPCR) and western blot analysis on brain lysates from embryonic day 14.5 (E14.5), postnatal day 0 (P0), and adult mice (>P56) showed that heterozygous mutations to *Chd8* resulted in decreased *Chd8* transcript and protein levels (Figure 1D-1E, S1C; Chd8 antibody: ab114126, Abcam). For the majority of the following studies, we analyzed mice harboring a 5 bp deletion in *Chd8* exon 5. Male *Chd8*^+/−^ mice were bred to wild-type females for at least four generations before further experiments, and multiple litters were used for all experiments to eliminate the potential impact of Cas9 off-target mutation. We tested for differences in brain size in *Chd8*^+/−^ mice at birth (postnatal day 0, P0), as macrocephaly is a hallmark trait in human *CHD8*^+/−^ individuals^8^. Maximal cortical anteroposterior length of *Chd8*^+/−^ brains was ~7% longer than matched wild-type (WT) littermates (ANOVA, p-value = 0.0440) with no significant differences between sexes (Figure 1F). These results show that *Chd8*^+/−^ mice are megalencephalic, suggesting neuropathological phenotypes that parallel *CHD8*^+/−^ humans.

**Figure 1.**
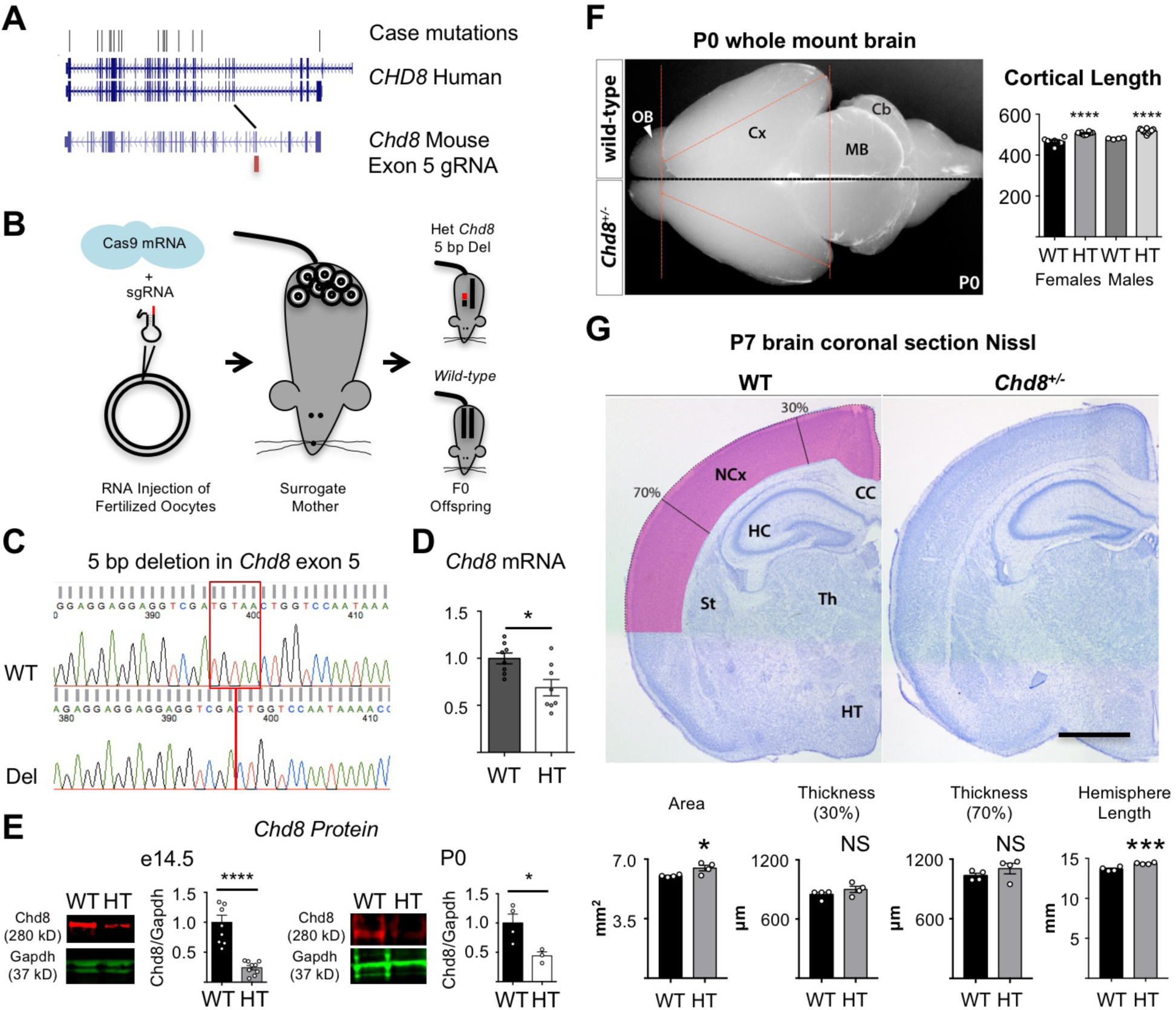
*Chd8*^+/−^ mouse model. **A**. Location of case mutations in human *CHD8* and corresponding guideRNA sequence homology for Cas9-targeting of mouse *Chd8*. **B**. Schematic of mouse line generation. **C**. Sequence trace showing 5 bp indel in exon 5. **D**. qPCR showing reduction of RNA in *Chd8*^+/−^ forebrain at P0 (n WT = 7, n *Chd8*^+/−^ = 5). **E**. Western blot of *Chd8*^+/−^ mice showing reduction of Chd8 protein (ab114126; Abcam) in *Chd8*^+/−^ forebrain at E14.5 (n WT = 9, *Chd8*^+/−^ = 9) and P0 (n WT = 4, *Chd8*^+/−^ = 3). **F.** Whole mount brain of *Chd8*^+/−^ mice at P0 reveals increased cortical length, indicative of megalencephaly. OB: olfactory bulb, Cx: cortex, MB: midbrain, Cb: cerebellum. WT n male = 10, female = 4; *Chd8*^+/−^ n male = 10, female = 4. **G.** *(Upper*) Representative coronal sections of wild-type and *Chd8*^+/−^ brains at P7 visualized with Nissl, n = 4 for both genotypes. Scale bar 1000 µm. *(Lower)* Plots (mean ± SEM with dots representing individual samples) of cortical area, thickness at 30% and 70% distance from the dorsal midline, and cortical hemispheric circumference. P-values derived using Student’s t-test for D, E and G and using ANOVA for F, *P < 0.05 ***P < 0.005 ****P < 0.001.

To further test the parameters of developmental megalencephaly in *Chd8*^+/−^ mutants, we examined brains in whole-mount and Nissl-stained coronal brain sections at P7 (Figure 1G), a time point after the conclusion of developmental neurogenesis and gliogenesis. At this stage, the mutant brains did not present overt neuropathological anomalies other than size increase. Total hemispheric circumference in *Chd8*^+/−^ brains was 4.7% longer (Student’s t-test, p = 0.0001) than in WT littermates (Figure 1G). We measured cortical thickness in two positions across the neocortex, at 30% and 70% distance from the dorsal midline. No significant differences between the genotypes were observed in either location (Student’s t-test, p = 0.2242; 0.2678), though *Chd8*^+/−^ mice trended larger across both measurements. Overall neocortical section area was ~8% higher in *Chd8*^+/−^ brains (Student’s t-test, p = 0.0009) compared to WT controls, confirming cerebral megalencephaly at this stage.

### Differential gene expression across neurodevelopment in Chd8^+/−^ mice

Having established neuroanatomical changes in *Chd8*^+/−^ mice that parallel clinical phenotypes described in *CHD8*^+/−^ humans and considering the role of *Chd8* in global transcriptional regulation, we profiled mRNA utilizing RNA-sequencing in forebrain dissected from four early developmental stages (embryonic days E12.5, E14.5, E17.5, and P0) and adult mice (>P56) (Figure 2A). After quality filtering, we individually analyzed 26 *Chd8*^+/−^ and 18 WT littermates, with full sample details reported in Table S1. Sample developmental stage represented the major components of gene expression variation, as expected given the large changes to transcription that occur across brain development (Figure S1). We observed reads overlapping the *Chd8* deletion sequence in all but one *Chd8*^+/−^ library and in no WT libraries. Decreased expression and corresponding decreased Chd8 protein levels were present in mice harboring the 5 bp exon 5 deletion as well as in the second line of mutant *Chd8*^+/−^ mice harboring an overlapping 14 bp deletion (Figure S1). For both mutant lines, *Chd8* deletion reads occurred at lower frequency relative to WT allele reads, suggesting that the frameshift transcript undergoes degradation (Figure S1).

**Figure 2.**
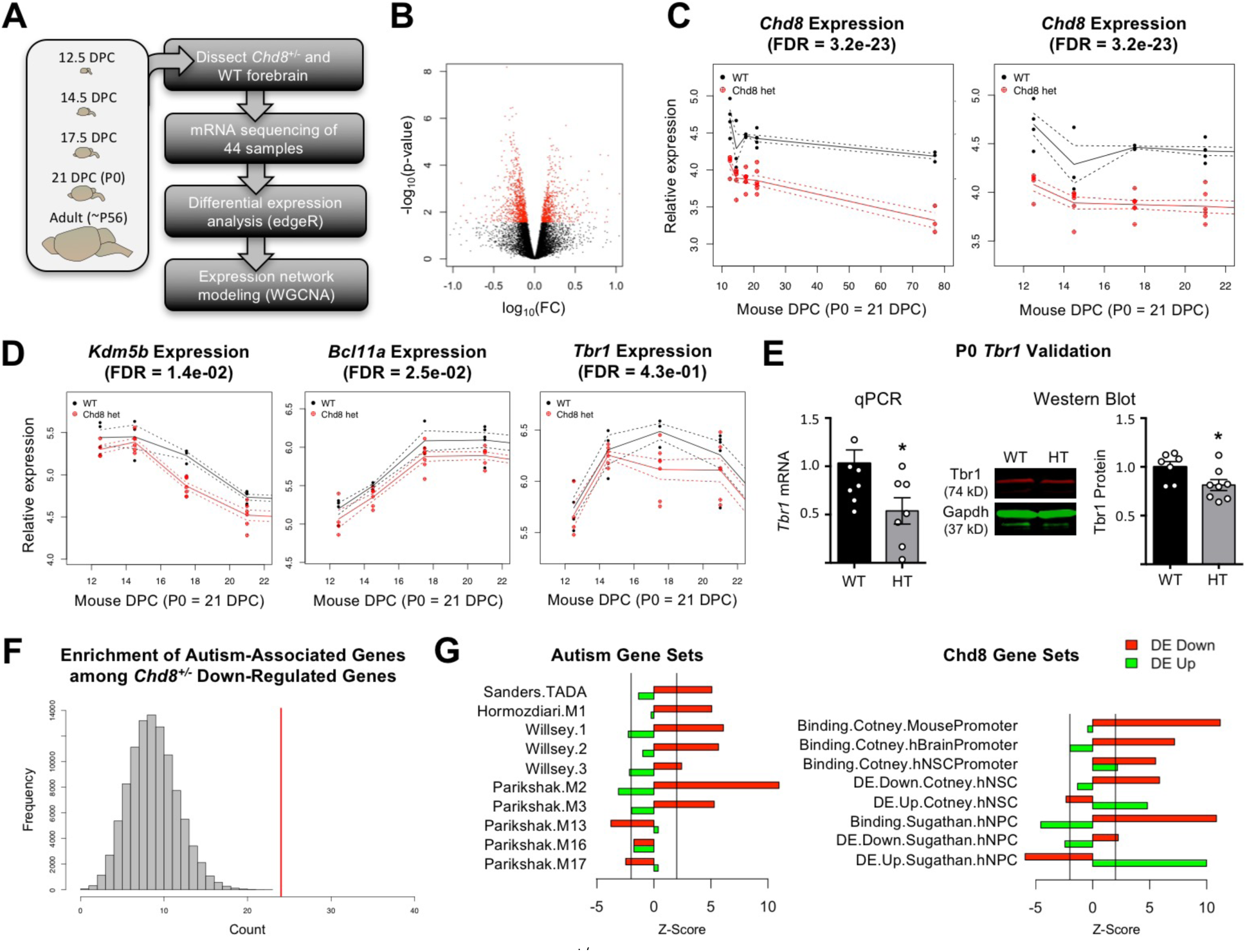
Differential gene expression in *Chd8*^+/−^ neurodevelopment. **A.** Schematic of our experimental pipeline: dissections of forebrain at five stages (E12.5, E14.5, E17.5, P0, ~P56) followed by RNA isolation, library preparation, sequencing and expression networking analysis (WGCNA). **B.** Volcano plot showing that most DE genes (plotted in red) exhibit relatively subtle fold changes. **C.** *Chd8* is the top differentially expressed gene, with panels showing expression log (RPKM) in *Chd8*^+/−^ and WT littermates across brain development (right panel shows only developmental stages). FDR of DE p-value shown. **D.** Example expression patterns in *Chd8*^+/−^ forebrain of three DE autism risk loci across developmental stages (FDR of DE p-value shown). **E**. Validation of DE expression of *Tbr1* RNA (left; n WT = 7, *Chd8*^+/−^ = 5) and protein (right; n WT = 7, *Chd8*^+/−^ = 8) in *Chd8*^+/−^ forebrain at P0 (Student’s t-test, *P = 0.0264, 0.0338). **G.** Distribution of expected number of autism-associated genes among based on 100,000 randomly sampled gene sets (grey bars) versus observed number of autism-associated DE genes in our dataset (red bar). **H.** Comparison of DE down- and up-regulated genes identified here with autism- and *Chd8*-binding genes. Z-score generated via permutation test comparing enrichment of test gene set to randomly sampled genes.

Using a statistical model that accounted for sex, developmental stage, and sequencing batch, we tested for differential expression across 14,163 genes that were robustly expressed in our datasets. At a significance cutoff corresponding to FDR < 0.05 (p-value < 0.0006), FDR < 0.1 (p-value < 0.0029) or FDR < 0.25 (p-value < 0.0272), 178, 418, and 1,536 genes, respectively, were differentially expressed (DE) (Table S2). While our full model is best suited for identification of genes where DE extends across developmental stages, we also examined stage-specific expression changes and full results are reported in Table S2. DE genes identified in our full model exhibited a range of expression trajectories across development, and the majority of significant expression changes in *Chd8*^+/−^ were relatively small (99.3% < 1.0 absolute log fold change, Figure 2B). These findings suggest that changes in neurodevelopmental gene expression are widespread yet subtle. We validated change in expression at P0 for a set of DE genes via qPCR (Figure S2, primers used reported in Table S3). Confirming our model, the top DE gene was *Chd8*, with decreased expression in the *Chd8*^+/−^ mice (log fold change = 0.59, p-value = 2.20E-27, FDR = 3.18E-23, Figure 2B). Irrespective of genotype, *Chd8* expression gradually declined across development in mouse forebrain, and significant reduction in expression was observed in *Chd8*^+/−^ mice at each stage (Figure 2C). No obvious isoform-specific changes in *Chd8* expression were present in the *Chd8*^+/−^ mice based on exon coverage (Figure S3). DE genes were significantly overrepresented among a number of Reactome^15^ pathways (Table S4). Example pathways with strong enrichment include RNA processing (e.g. Processing of Capped Intron-Containing Pre-mRNA, FDR = 1.59E-09), gene expression (FDR = 1.66E-04), cell cycle (FDR = 0.002), Regulation of TP53 Activity (FDR = 0.009), and Axon Guidance (FDR = 0.009). Similar overall pathway enrichment was observed for DE genes at FDR cutoff of 0.05, 0.1, and 0.25. The strong signature of differential expression enabled us to map the perturbation of biological pathways and processes caused by *Chd8* haploinsufficiency across neurodevelopment.

### Overlap of autism-relevant genes and Chd8 binding targets with Chd8^+/−^ DE genes

First we examined whether *Chd8* acts as a regulator of autism-linked genes during brain development. In agreement with previous *in vitro* knockdown models^12,14^, autism risk genes were among DE genes in *Chd8*^+/−^. Figure 2D shows example DE high-confidence autism risk loci (*Kdm5b*, *Bcl11a*, and *Tbr1*) with different developmental expression patterns. We validated decrease in mRNA expression and protein level for *Tbr1* at P0 in *Chd8*^+/−^ forebrain via qPCR analysis and western blot (Figure 2E, Figure S4). We next tested for overlap between DE down-regulated and up-regulated genes (FDR < 0.25) and published gene sets of relevance to autism genetics and *Chd8* regulation. For example, of the 143 genes implicated by presence of mutations in autism cases^16^ that were detected in our expression data, 24 were DE at FDR < 0.25 and down-regulated (permutation test p-value = 3.08E−07) (Figure 2F). We also observed significant enrichment among down-regulated DE genes with autism risk genes identified by other studies^13,17^ (Figure 2G). We examined global gene co-expression networks relevant to autism as identified via network analysis of human neurodevelopmental gene expression^18^, including two early developmental autism-associated networks (Parikshak.M2 and Parikshak.M3) as well as three autism-relevant networks expressed later in human brain development (Parikshak.M13, Parikshak.M16, Parikshak.M17). We observed strong enrichment specific to the early developmental networks among the global set of *Chd8*^+/−^ down-regulated genes (Parikshak.M2 p-value = 5.30E-28; Parikshak.M3 p-value = 1.20E-07). We observed no global enrichment among DE down-regulated genes for the later developmental modules or among FMRP targets^19^ and gene networks identified in postmortem autism case brains^20^.

Next we asked whether our DE data is consistent with CHD8 binding and differential expression after *CHD8* knockdown in human *in vitro* models, as reported in previous studies^12,14^. While these studies show that CHD8 directly regulates a large number of promoters across mouse and human systems, we nonetheless observed consistent enrichment among down-regulated DE genes in *Chd8*^+/−^ forebrain for CHD8 target genes identified in these earlier studies (Figure 2G). For example, down-regulated DE genes from our study were enriched for genes with Chd8 promoter binding in E17.5 mouse frontal or occipital cortex^14^ (p-value = 4.70E-29). There was no enrichment among our up-regulated DE sets for genes targeted by CHD8, suggesting that up-regulation is indirect or occurs at earlier time points. Comparing DE genes that exhibited down- or up-regulation in our study with genes that show DE in the matched direction in the independent knockdown studies discussed above, we observed strong direction-specific enrichment between our study and these two previous studies (Figure 2G).

Finally, we see that differential expression is not limited to early developmental effects. For example, many synaptic genes are also impacted (Figure S5). Consistent with previous studies, we observed that Chd8 regulates TP53 and, to a lesser degree, *Wnt* signaling pathways that control processes from early neurodevelopment to synaptic function^10,21,22^. As such, *CHD8* haploinsufficiency may drive autism-associated pathology via multiple neurodevelopmental mechanisms. This analysis confirms that Chd8 is required either directly or indirectly for typical expression of autism-relevant gene networks during neurodevelopment.

### Gene expression network analysis to identify perturbations to Chd8^+/−^ neurodevelopment

We next explored how DE genes are organized into networks that follow parallel expression trajectories during brain development with a goal of identifying stage-specific neurodevelopmental processes that are perturbed in *Chd8*^+/−^ mice. We used weighted gene correlation network analysis (WGCNA^23^) to identify co-regulated gene modules from our developmental transcriptomic data. Fifteen discrete modules were identified that exhibited specific trajectories of expression and covariation across forebrain development (Figure 3A-3B, Figure S6, Table S5). DE genes assigned to specific modules were enriched for relevant gene sets (Figure 3C) and stage-specific Gene Ontology Biological Process annotation terms (Figure 3D, Table S7), confirming that the modules captured by this approach are developmentally and biologically relevant.

**Figure 3.**
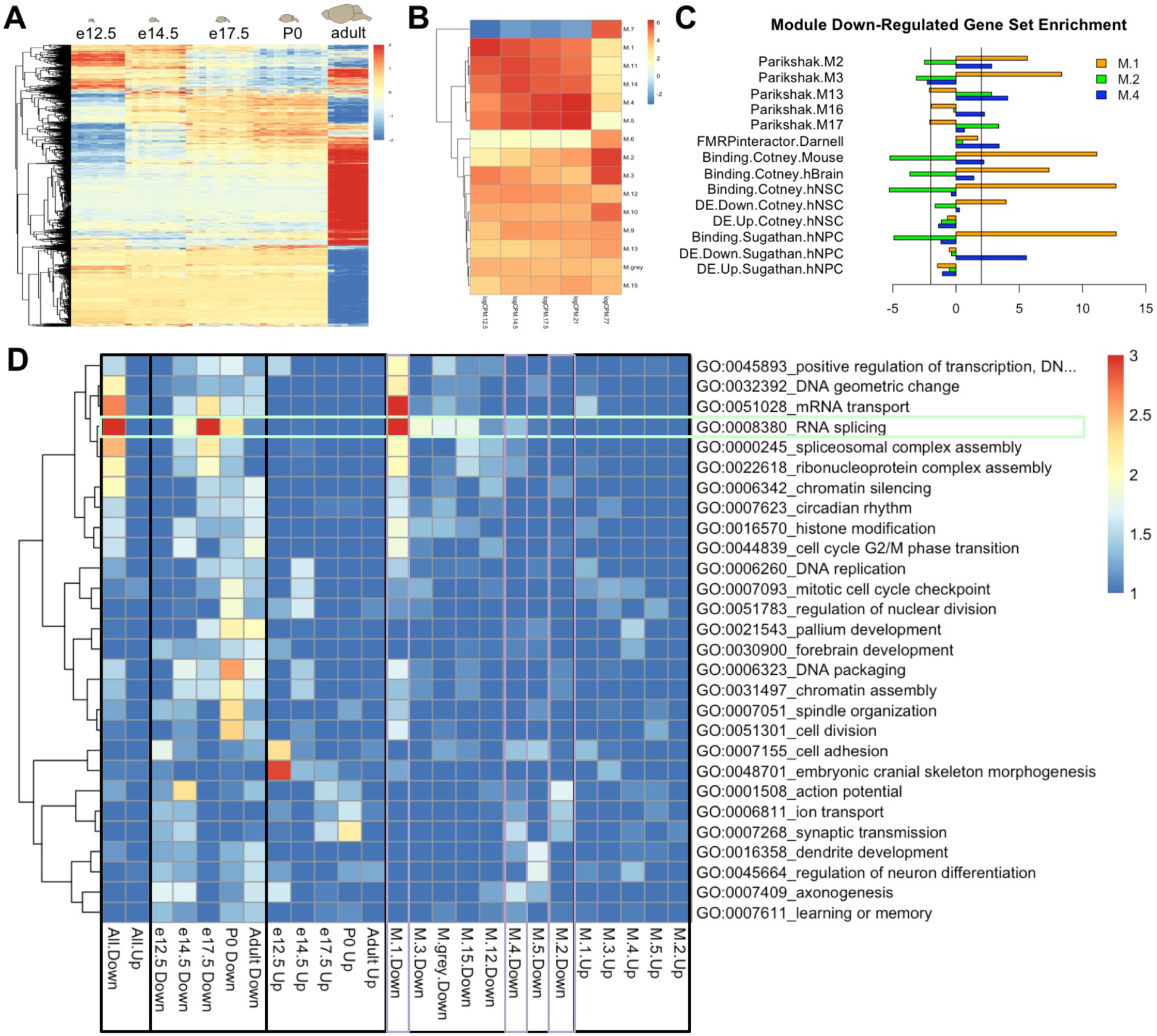
Identification of DE genes with correlated expression patterns across brain development reveals perturbation to early and later neurodevelopmental pathways. **A.** Heatmap representing expression of DE genes across all samples and stages. **B.** Mean expression across developmental stages for the 15 developmental gene expression modules. **C.** Permutation test of overlap between module-specific DE down-regulated genes from M.1 (early neurodevelopment), M.2 (neuronal differentiation), and M.4 (late neuronal/brain genes) (purple boxes in panel D) and autism and *Chd8*-relevant genes sets. M.1 DE down-regulated genes are enriched for early expressed autism-associated networks (Parikshak M2 and M3) and direct binding by *Chd8*, while M.2 and M.4 are enriched for FMRP targets and later autism-associated gene networks (Parikshak M13, M16, M17) but not Chd8-bound genes or early developmental networks. **D.** Functional enrichment of GO biological process annotations for stage- and module-specific DE genes.

First, we looked at module representation among high-confidence autism genes identified in autism genome sequencing efforts^16^. While these genes were strongly enriched overall among DE down-regulated genes (Figure 2F), autism risk genes are present across modules rather than exhibiting strong module-specific clustering. For example, M.1 is characterized by decreasing expression across neurodevelopment, is associated with chromatin, RNA processing, and cell cycle, and includes the largest number of DE down-regulated autism risk genes (e.g. *Kdm5b* and *Chd2*). M.1 is discussed in more detail below. M.4 represents a network of genes with rising expression from E12.5 to P0 with much lower expression in adult brain. Down-regulated genes in M.4 are enriched for GO terms associated with transient neuronal differentiation processes (e.g. axonogenesis) and include autism risk genes involved in post-mitotic migration and neuronal maturation (e.g. *Bcl11a*, *Ank2,* and *Ctnnbp2)*. M.2 is characterized by low expression early that gradually increases at each stage including in adult forebrain. Down-regulated genes in M.2 are enriched for GO terms such as synaptic transmission that are hallmarks of more mature neurons, and include autism risk genes such as *Cers4* and *Gria1*. Many synaptic genes are down-regulated in *Chd8*^+/−^ forebrain, with down-regulation strongest in the adult time point but often present developmentally as well (Figure S5), including high-confidence autism risk genes such as *Scn2a1* (M.5) and *Cacna1b* (M.2). Segregation of DE genes into modules revealed Chd8 haploinsufficiency causes perturbation of autism-relevant gene sets that appeared to be in separate causal pathways^6^. This examination suggests a developmental hierarchy of autism-relevant pathology in *Chd8*^+/−^ mice involving chromatin remodeling, transcriptional regulation, and synapse function, the three major pathological pathways implicated via human genetics^6^.

In contrast to the spread of high confidence autism risk genes across developmental modules, we observed strong module-specific enrichment for autism-associated developmental expression networks identified by Parikshak et al. 2013, as well as for CHD8 binding and knockdown data, and DE genes whose transcripts are FMRP targets (Table S6). To illustrate this, we highlighted gene set enrichment for down-regulated DE genes from three modules with distinct developmental trajectories discussed above, M.1, M.4, and M.2 (Figure 3C). We observed enrichment in overlap with genes in Parikshak.M3, the earliest expressed network identified in Parikshak et al. 2013 that was specific to down-regulated genes in M.1, our early expression module. Genes in Parikshak.M2, a module that is anchored later than Parikshak.M3 but still representative of early brain development, overlap both M.1 and M.4. The three late modules identified by Parikshak et al. 2013 overlap with down-regulated genes from either M.4 (Parikshak.M13 and Parikshak.M16) or M.2 (Parikshak.M17), but not M.1. While we did not observe global enrichment among all DE genes for FMRP targets^19^, there was module-specific enrichment for M.4 DE down-regulated genes among FMRP targets. We observed strong module-specific enrichment of M.1 for Chd8 binding targets identified in embryonic brain and in vitro models of neural develoment^12,14^, suggesting direct regulation of DE genes by Chd8. Finally, we found module-specific enrichment of genes down-regulated after CHD8 knockdown in hNSC (M.1) and hNPC (M.4), suggesting similar gene regulatory consequences produced by *Chd8* haploinsufficiency across systems.

### Chd8^+/−^ mice exhibit down-regulation of genes involved in RNA processing in forebrain across developmental expression modules

A number of GO terms were strongly enriched for DE genes mapping across modules, suggesting perturbation to stage-specific gene sets controlling these processes across developmental stages (Figure 3D). This signature was particularly strong for genes involved in RNA processing (Figure 4). DE genes were significantly overrepresented among genes annotated to RNA processing, e.g. Processing of Capped Intron-Containing pre-mRNA (FDR = 1.59E-09), mRNA Splicing (FDR = 2.90E-09), and mRNA 3’-end Processing (FDR = 3.29E-04) in the Reactome database (Figure 4A, Table S4). These genes represented modules with highly divergent expression trajectories (Figure 4B). For example, among genes annotated to RNA splicing, expression of *Dhx9* (M.1) decreases across neurodevelopment and has not been functionally characterized in brain but has been reported in autism-risk networks^17^ (Figure 4B), while *Upf3b* (M.15) expression increases across development and is a neuron-specific factor required during neuronal differentiation that is implicated in intellectual disability^24,25^ (Figure 4B). While RNA processing genes present different overall expression trajectories across brain development, perturbation to expression generally peaked at E17.5 for these genes, suggesting that this represents a period where this process is critical in brain development. Unlike synaptic genes and critical transcription factor genes (e.g. *Tbr1*), the DE genes annotated to have a role in RNA processing have not been well studied in the context of brain development and represent candidates for future investigation. RNA processing genes that are DE in *Chd8*^+/−^ mice are reported in Table S8.

**Figure 4.**
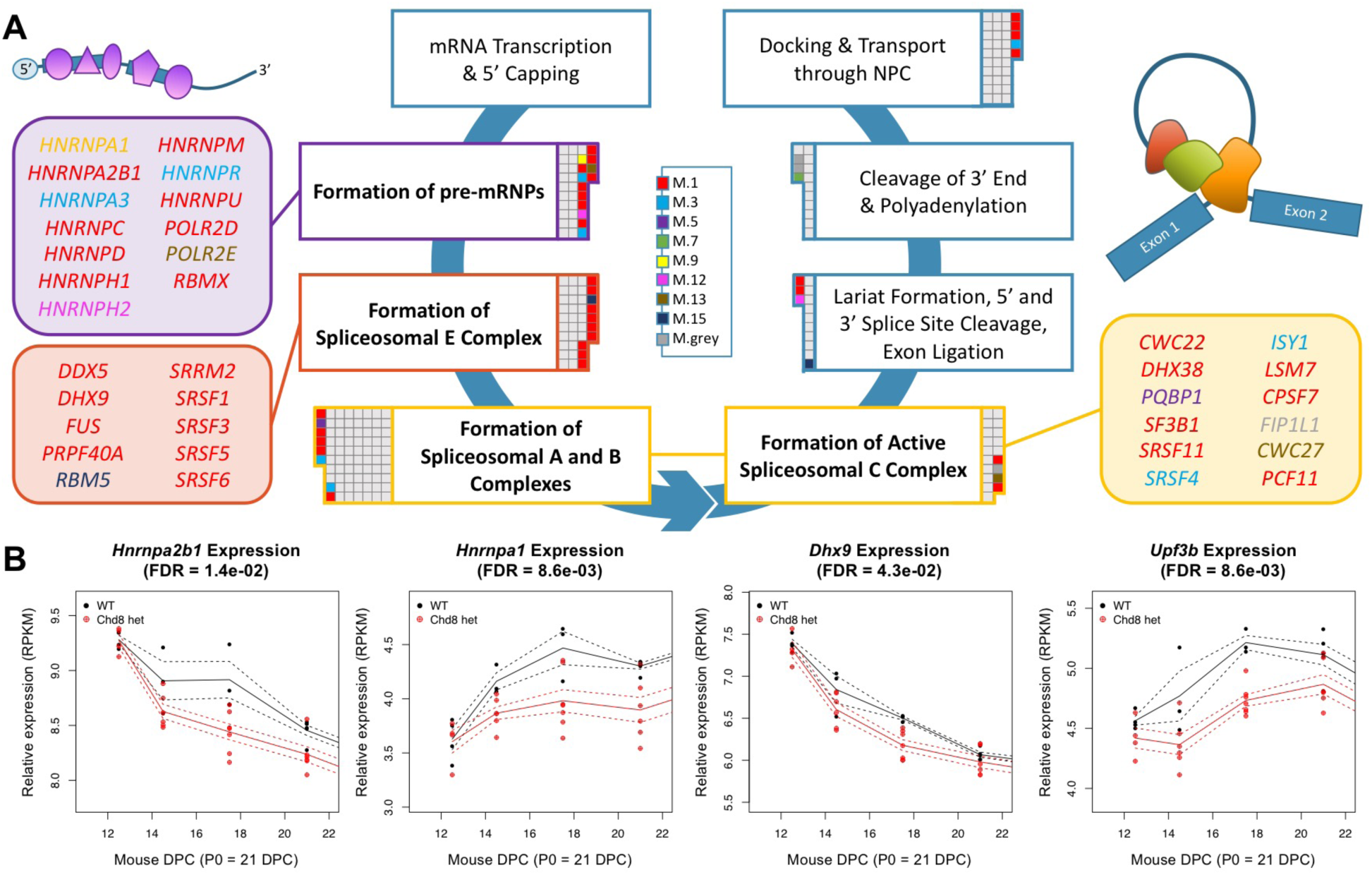
RNA processing pathways are enriched for differentially expressed genes in *Chd8*^+/−^ mice. **A.** Condensed model of the RNA processing pathway on Reactome, adapted from the parent pathway *Processing of Capped Intron-Containing pre-mRNA*. Genes annotated to different steps in the RNA processing pathway are denoted as boxes. DE genes are colored by module membership; non-differentially expressed genes are white. Each gene is represented once, at the first step the gene appears in the pathway. **B.** Developmental expression trajectories of example DE down-regulated genes annotated to the formation of pre-mRNPs (*Hnrnpa2b1*, *Hnrnpa1*), formation of the spliceosomal E complex (*Dhx9*), and lariat formation and 5’ splice site cleavage (*Upf3b*). DE p-value FDR shown.

### Perturbation to early developmental expression network and associated increase in neuronal proliferation in Chd8^+/−^ mice

M.1 consists of 3,590 genes that show a general trend of decreasing expression levels across neurodevelopment (Figure 5A). M.1 showed the strongest enrichment for autism risk early developmental modules^18^ and for Chd8 binding targets in embryonic brain and in vitro models. M.1 also had the greatest enrichment of down-regulated genes (p-value = 5.39E-21). 454 genes in M.1 are DE at FDR < 0.25 (350 down-regulated, 104 up-regulated), accounting for ~30% of all DE genes identified in our study. We examined change in expression of DE genes within M.1 at each developmental stage, observing that up-regulation peaked at E14.5 compared to a fold change peak for down-regulated genes at E17.5 (Figure 5B). Analysis of protein-protein interactions (STRING^26^) showed that DE genes in M.1 had more protein-protein interactions than expected by chance (observed edges = 972, expected edges = 237, enrichment = 4.10, STRING p-value < 0.0001). Interacting genes in M.1 were enriched for functions regulating genome structure, cellular proliferation and differentiation, with enrichment for GO terms (Table S9) including RNA processing, chromatin organization, and mitotic cell cycle (Figure 5C). M.1 DE genes annotated to these terms, including a number of autism risk genes, were identified as Chd8 targets. A subset of these genes was also DE after *CHD8 in vitro* knockdown in human cells (Figure 4D). This analysis suggests that Chd8 binding directly regulates M.1 genes and that differential expression of M.1 genes in *Chd8*^+/−^ embryos may drive changes in chromatin structure and RNA metabolism linked to early neurodevelopmental pathology associated with disruption to proliferation and neuronal differentiation.

**Figure 5.**
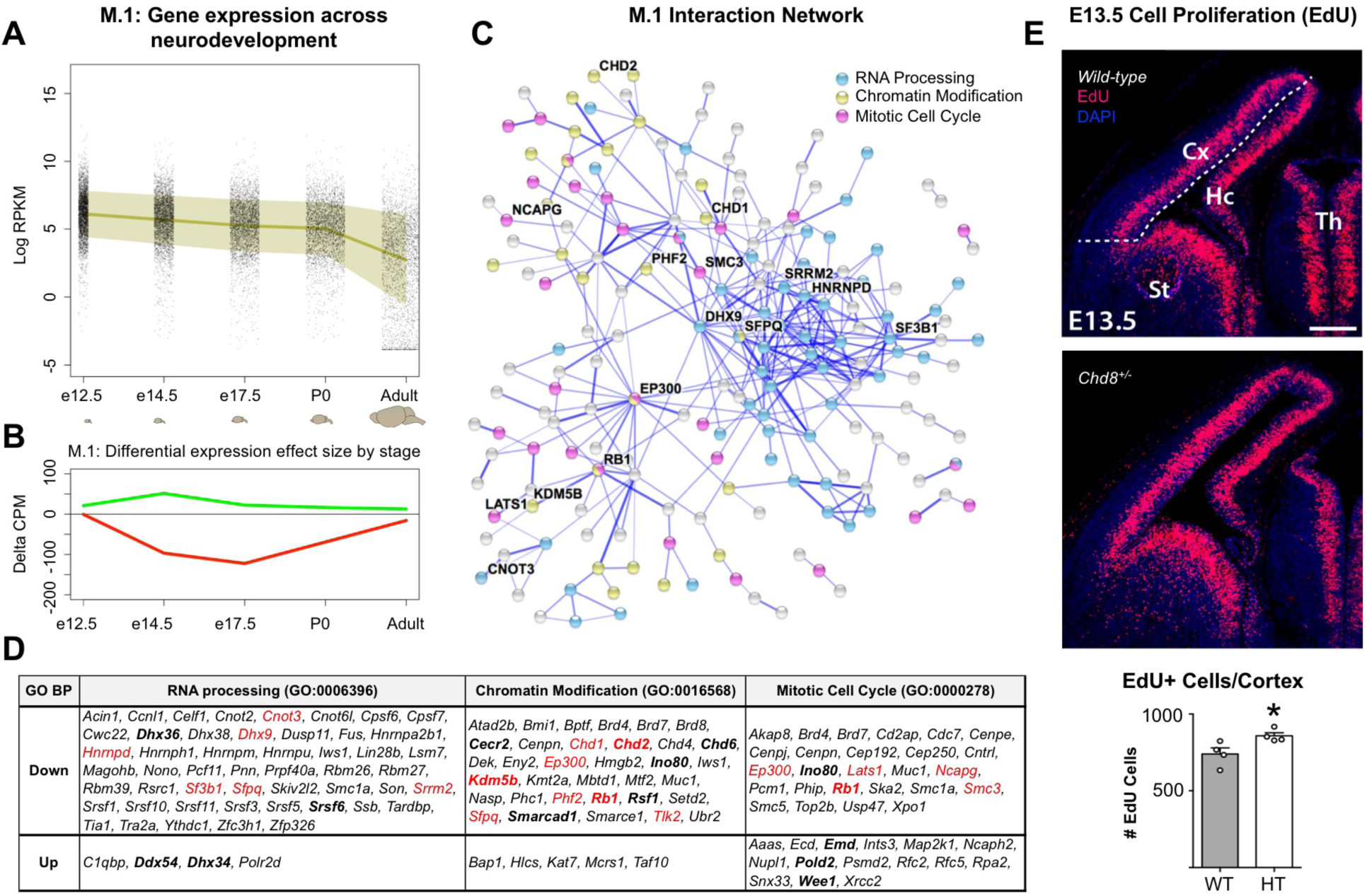
An early neurodevelopmental expression network (M.1) regulated by Chd8 haploinsufficiency involved in chromatin organization, RNA processing, and cell cycle. **A.** M.1 gene expression plotted across brain development. Dots represent individual genes, line represents mean expression and shaded area represents ± 1 SD. **B.** Relative mean differential expression of up-regulated (green) and down-regulated (red) genes in M.1 across brain development. **C.** Protein-protein interaction network of M.1. DE genes are colored by annotation to chromatin organization, RNA processing, and mitotic cell cycle GO Biological Process terms. Labeled genes have been previously identified as autism risk genes. **D.** M.1 Chd8-bound DE genes associated with selected GO terms. Red: autism risk genes; black: differential expression in previous *in vitro* Chd8 knockdown studies. **E.**(*Upper*) Coronal section of E14.5 stained for EdU (m*agenta*), a marker of proliferation, and DAPI (*blue*) in WT and *Chd8*^+/−^ mice (n = 4 for both genotypes). Scale Bar 200 µm. (*Lower*) Plot (mean ± SEM with dots representing individual samples) of EdU positive cells/area. Student’s t-test p-value = 0.0338.

**Figure 6.**
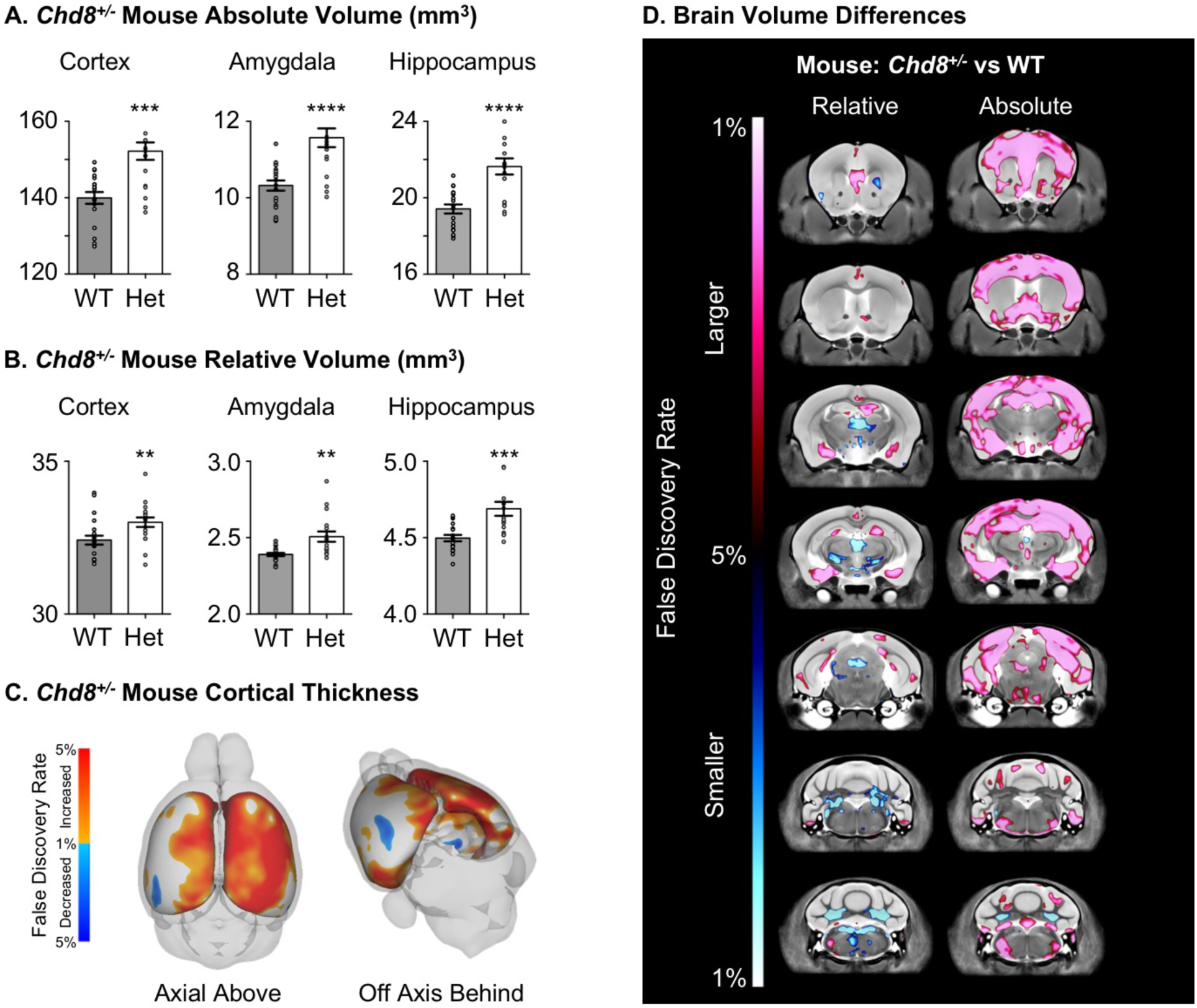
*Chd8* haploinsufficiency drives macrocephaly and cognitive impairment in mouse and human. **A.** MRI revealed significant increases in absolute regional volume of cerebral cortex, amygdala, and hippocampus (****p < 0.0001) in *Chd8*^+/−^ mice. N: WT=21, *Chd8*^+/−^=18. See Table S11 for full regional statistical analysis and FDR values. **B.** Increases in volume are still significant for the cortex (**p = 0.0097), amygdala (**p = 0.0014) and hippocampus (***p = 0.0004) after correction for absolute brain volume. (Student’s t test p-value, error bars denote SEM). **C.** Increased cortical thickness is present in *Chd8*^+/−^ mice. **D.** Voxel-wise differences in volume between *Chd8*^+/−^ and WT littermates.

To examine whether these alterations in developmental genes play a functional role in neuronal development that could lead to megalencephaly, we performed 5-ethynyl-2′-deoxyuridine (EdU) proliferation assays (Figure 5E). Assessing the number of EdU^+^ cells in the cortical ventricular zone (VZ) after a 1.5 hour pulse at E13.5, a time point of peak neurogenesis, we found their number significantly increased in the mutant by 15.9% (Student’s t-test, p = 0.0338, n = 4 for either genotype). Considering that a number of genes associated with brain development and cortical structure exhibited DE, we also examined cortical morphology via analysis of Tbr1 and Ctip2 immunostaining at P0 (Figure S7). We observed no gross alterations to lamination and found no evidence for focal cortical lesions (n: WT = 8; *Chd8*^+/−^ = 10). These findings experimentally corroborate altered proliferative dynamics in *Chd8*^+/−^ mutants, linking altered neurogenesis and megalencephaly in *Chd8*^+/−^ mice.

### Analysis of Chd8^+/−^ adult brain structure via MRI

To establish whether structural changes persist in the *Chd8*^+/−^ mouse brain, structural MRI was performed to identify changes in regional brain volume and connectivity between adult *Chd8*^+/−^ mice (n=18) and matched wild-type littermates (n=21). No significant differences were observed in body weight or other relevant measures of general health in adult *Chd8*^+/−^ mice (Table S10). Neuroanatomy was assessed and volume was measured as absolute volume (mm^3^) and relative volume (% total brain volume). Considering regional differences, the most affected region was the cortex, which was increased by 7.5% with a false discovery rate (FDR) of 1%. Similarly, the cerebral white matter and cerebral gray matter were also larger in the *Chd8*^+/−^ mice at 5.4% (FDR of 3%) and 6.1% (FDR of 2%), respectively. When the male and female mice were examined independently, female *Chd8*^+/−^ mice exhibited stronger effect sizes but both sexes exhibited overall similar trends. In addition to the summary regions, 159 independent brain regions were assessed with divisions across the cortex, subcortical areas, and cerebellum. Full results for comparison across individual brain regions are reported in Table S11. *Chd8*^+/−^ mice showed robust increase in absolute volume across cortical regions, hippocampus (+10.3%, FDR < 1%), and amygdala (+11.0%, FDR < 1%) (Figure 5A-5B). The *Chd8*^+/−^ mice also displayed increased cortical thickness, particularly along the cingulate cortex (Figure 5C). After correction for total volume, relative volumes were still significantly larger, though cortex failed to surpass the FDR < 5% cutoff. Deep cerebellar nuclei showed decreased relative volume (-1-3%, FDR < 2%). Voxel-wise differences showed similar trends (Figure 5D). Diffusion Tensor Imaging (DTI) revealed no differences in fractional anisotropy or mean diffusivity in either the regional or voxel-wise measurements, indicating that the anatomical connectivity of the white matter in the *Chd8*^+/−^ mice was not significantly different from WT littermates (not shown).

### Behavioral phenotyping of adult Chd8^+/−^ mice

Behaviors relevant to ASD were assessed in adult *Chd8*^+/−^ mice using two assays of social behaviors and two assays of repetitive behaviors (Figure 7), as previously described^27^. In the 3-chambered social approach test^28^, normal sociability was detected in both genotypes (Figure 7A-7B). Time spent in the chamber with the novel mouse was greater than time spent in the chamber with the novel object, meeting the definition of sociability in this assay, for both WT and *Chd8*^+/−^ (Figure 7A; WT: t (1, 40) = 6.07, p < 0.001; *Chd8*^+/−^: t (1,34) = −3.93, p < 0.001). No sex differences were detected (F (1, 37) = 2.16, p > 0.05). Time spent sniffing the novel mouse was greater than time spent sniffing the novel object in both WT and *Chd8*^+/−^ (Figure 7B; WT: t (1, 40) = 2.47, p < 0.02; *Chd8*^+/−^: t (1, 34) = −2.33, p < 0.03) again without sex effects (F (1, 37) = 0.107, p > 0.05). Number of entries into the side chambers was not affected by genotype in the social phase (Figure 7C; F (1, 37) = 0.99, p > 0.05), nor on the entries into the left or right chambers parameter during the previous habituation phase (F (1, 37) = 0.069, p > 0.05), indicating normal general exploratory activity in both genotypes during the social approach assay.

**Figure 7.**
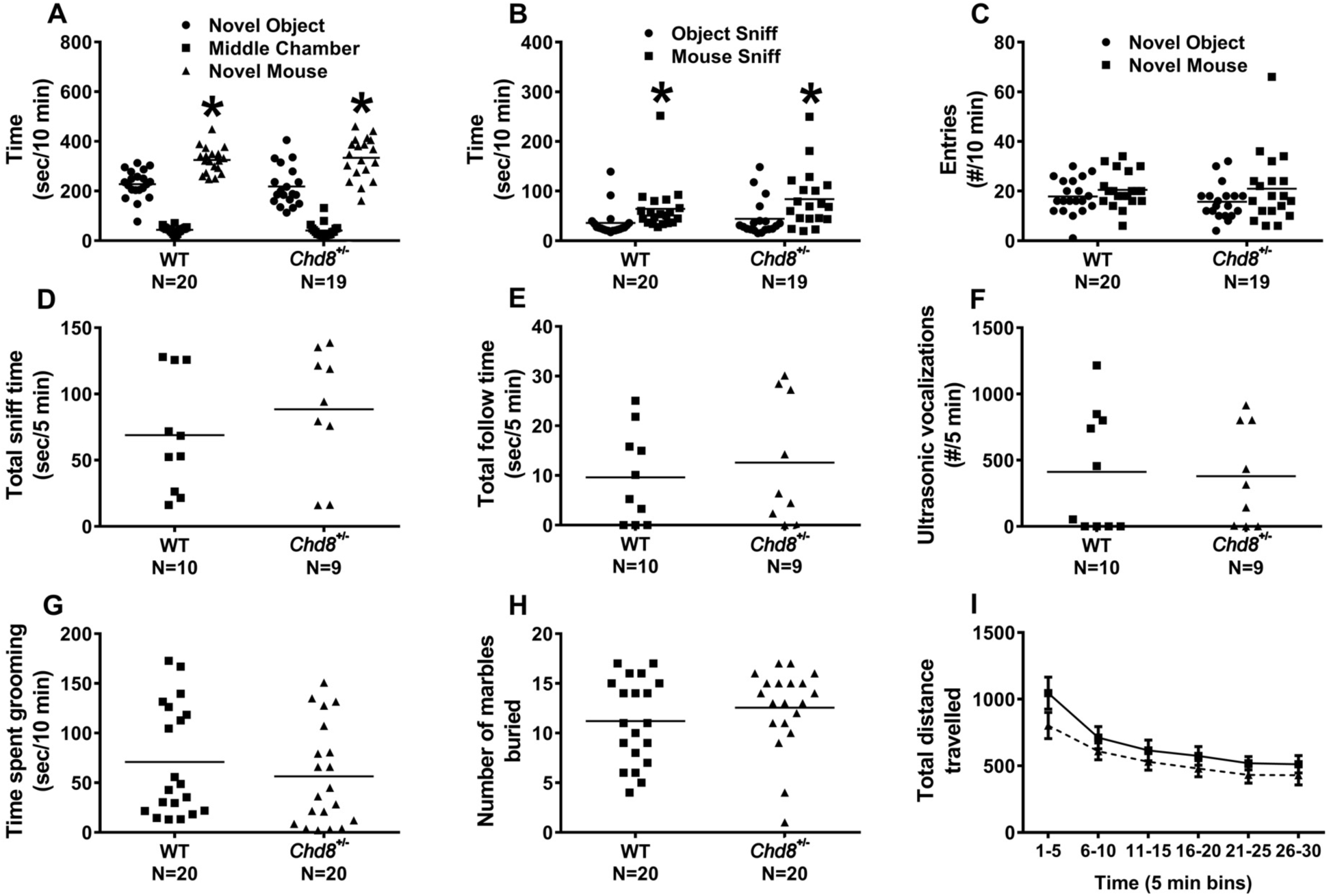
Adult *Chd8*^+/−^ mice do not differ from WT on ASD-relevant social and repetitive behavior assays. *Chd8*^+/−^ mice and WT littermate controls both met the definition of sociability on the 3-chambered social approach test. (**A-C**). Each genotype spent significantly more time in the chamber with the novel mouse than in the chamber with the novel object, spent significantly more time sniffing the novel mouse than sniffing the novel object, and showed normal locomotor entries between chamber and normal male-female social interactions with concomitant ultrasonic vocalizations (**D-F**). *Chd8*^+/−^ and WT displayed similar amounts of self-grooming (**G**), marble burying (**H**), and open field exploratory locomotion (**I**).

No abnormalities were observed on social parameters in male *Chd8*^+/−^ during the male-female reciprocal social interaction test, in which male WT and *Chd8*^+/−^ subject mice were individually paired with an unfamiliar estrous B6 female (Figure 7D-7E). WT and *Chd8*^+/−^ males spent similar amounts of time engaged in sniffing (Figure 7D; t (1, 17) = 0.35), and following (Figure 7E; t (1, 17) = 0.78, p > 0.05) the female, with numerical scores comparable to previous findings^29,30^. Ultrasonic vocalizations recorded during male-female interaction showed no genotype difference in the number of emitted calls (Figure 7F; t (1, 17) = −0.74, p > 0.05).

No spontaneous stereotypies or repetitive behaviors were observed. Genotypes did not differ on time spent in self-grooming behavior (Figure 7G; t (1, 38) = −1.29, p > 0.05), or in numbers of marbles buried (Figure 7H; t (1, 38) = 0.63, p > 0.05). No sex differences were detected (self-groom: t (1, 37) = −1.79, p > 0.05; marble burying: t (1, 37) = −1.49, p > 0.05). Open field locomotor activity did not differ between genotypes (Figure 7I, F(1,27) = 0.28, p>0.05), and both genotypes displayed the expected habituation to the novel environment across the 30 minute test session (Figure 7I; F (5, 90) = 10.62, p < 0.001), indicating normal exploratory behavior.

Taken together, these results indicate that heterozygous loss of *Chd8* did not significantly alter social approach, reciprocal social interaction, or repetitive behaviors in this first cohort of adult *Chd8*^+/−^ mice, as conducted with established assays with face validity to the diagnostic symptoms of ASD.

## Discussion

This integrative analysis of a new mouse model demonstrates that *Chd8* haploinsufficiency results in ASD-relevant neurodevelopmental phenotypes. Our results show that germline 5 bp and 14 bp deletion mutations in *Chd8* exon 5 result in decreased mRNA, consistent between qPCR and RNA-seq. *Chd8* itself was the most strongly significant DE gene of our transcriptomic analysis, and was DE at all developmental stages. Further, decrease in allele-specific expression of the mutant allele was evident in RNA reads. Such reads occur at low levels relative to the WT allele, suggesting degradation of the mutant frameshift transcript via nonsense-mediated decay. Finally, the amount of Chd8 protein assayed via western blot was consistently decreased in *Chd8*^+/−^ mouse brain across developmental stages. These results indicate that the short deletions in *Chd8* exon 5 in our mouse models result in a germline haploinsufficient state, with concomitant decrease in *Chd8* mRNA and protein observed in forebrain across all methods and experiments.

Our experiments confirm that *Chd8* regulates proliferation and neurogenesis, and suggest substantial impact across neurodevelopment. They also suggest similar binding targets and biological function in mouse and human neurodevelopment and a parallel causal role of *CHD8* haploinsufficiency in megalencephaly. We observed significant overlap between down-regulated genes in *Chd8*^+/−^ mice and autism risk gene sets produced by independent groups using a variety of approaches^13,16-18^. This overlap suggests that perturbation to neurodevelopment in *Chd8*^+/−^ mice parallels autism-relevant human neurobiology, a finding consistent with our neuroanatomical and structural MRI results.

Our RNA analysis captures subtle changes in transcription across brain development in *Chd8*^+/−^ mice. These changes were consistent across developmental stages for perturbed genes, were highly relevant to ASD-associated networks, and strongly correlated with biological pathways and expression modules of interest. Our results parallel *in vitro* findings that suggested convergence across risk pathways after CHD8 knockdown^12^, providing a developmental framework revealing disruption of convergent ASD pathways in a genetic mouse model of Chd8 haploinsufficiency. Unlike genomic analysis using *in vitro* knockdown studies of *CHD8*^12,14^, our network analysis using *in vivo* data enabled characterization of the impact of *Chd8* haploinsufficiency across neurodevelopment.

Our results suggest a developmental hierarchy of changes in *Chd8*^+/−^ brain development. For example, M.1, the module with the strongest enrichment for DE genes and genes directly targeted by CHD8, represents a highly interactive network of genes central to control of chromatin state and RNA processing, including genes implicated in autism. M.1 may represent a critical network directly regulated by *Chd8*. The timing disruption of this module supports that it plays a central role in the neuronal proliferation phenotype observed in E13.5 *Chd8*^+/−^ brain and may be functionally linked to the impact on RNA processing gene expression. The overlap of RNA processing gene down-regulation in M.1 and in later developmental expression modules suggests that *Chd8* haploinsufficiency results in general changes in molecules involved in RNA processing during neuronal differentiation. Considering the large number of DE genes involved in RNA processing, our results further indicate that disruption to RNA processing is an important player in neurodevelopmental disorders, in line with other neurodevelopmental disorder genetic models such as *Fmr1*^31^ and *Rbfox1*^32^. Furthermore, though alternative splicing has been well-established to a critical role in cell differentiation and proliferation in the developing brain^33^, the roles of other RNA processing pathways such as RNA stability or translocation in brain development are less studied^34^ and could provide potential candidates for future investigation.

While dysregulation of genes in M.1 and of RNA processing genes peaks at E17.5, our analysis suggests changes across neurodevelopment driven by *Chd8* haploinsufficiency. These changes indicate convergent neuropathology connecting chromatin remodeling, neuronal differentiation, and synaptic pathways, the principle gene networks identified in autism case sequencing studies. For example, we observed down-regulation of genes involved in RNA processing (e.g. *Upf3b* and *Hnrnpd*), neuronal differentiation (e.g. *Bcl11a* and *Tbr1*) and in synapse development and function (e.g. *Scn2a1* and *Cacna1b*), all of which have been implicated in autism via human genetic studies. The transcription data generated for this study will be a useful resource for future dissection of pathways involved in the pathogenesis of neurodevelopmental disorders and in classification of risk genes from genetic studies. Further studies may capture the neuroanatomical and cellular changes and perturbed signaling pathways associated with differential expression signatures in *Chd8*^+/−^ brain development. Future work will also be necessary to determine the stage- and cell-specific role of Chd8-binding to establish and maintain expression patterns of these genes.

Structural changes in the brain of adult *Chd8*^+/−^ mice parallel other relevant mouse models. A recent study examined 26 different mouse models related to autism^35^, clustering these models into 3 distinct groups. Key aspects of Group 1 included larger sizes of cortical structures, particularly the frontal and parietal lobes, and smaller sizes in the cerebellum, which is in line with the *Chd8*^+/−^ mouse described here. This group of models included *Nrxn1α*, *Shank3, En2*, and *Fmr1*. The *Chd8*^+/−^ mouse most resembled the differences found in the *Fmr1* mutant mice. Further examination may reveal similarities with other mouse models within this group beyond neuroanatomy (e.g. excitatory deficits in the *Nrxn1α* mouse^36^), as suggested by the widespread transcriptional changes present in *Chd8*^+/−^ neurodevelopment. Increases in cortical anteroposterior length and developmental neurogenesis appear largely overlapping in *Chd8*^+/−^ mice and *Wdfy3* mutants, a recently reported model of megalencephaly in autism^37^.

In comparison to the *Chd8*^+/−^ mice studied here, heterozygous mouse models of *Pten*, another gene associated with ASD and macrocephaly, do exhibit core aspects of ASD^38^. However, studies of other mouse models of genes implicated in ASD have not identified behavioral phenotypes with face value to ASD. Additional studies are necessary to further examine behavioral phenotypes at earlier developmental stages to test for causal relationship between structural changes and behavior in *Chd8*^+/−^ mice. The presence of genomic and neuroanatomical phenotypes in *Chd8*^+/−^ mice that parallel the clinical signature of human *CHD8* mutations suggests similar neurodevelopmental pathology between human and mouse. Intriguingly normal phenotypes on the autism-relevant social and repetitive assays conducted here highlight future opportunities for comprehensive behavioral phenotyping in a replication cohort, to evaluate social and repetitive behaviors at juvenile ages and to investigate phenotypes relevant to other symptom domains described for individuals with *CHD8* mutations, including cognitive impairments and attentional disorders.

Our initial survey of mice heterozygous for mutation to *Chd8* revealed significant findings across genomic and anatomical axes of neurobiology. These experiments link increased regional brain volume to perturbations of biological pathways across neurodevelopment, recapitulating primary neuroanatomical traits observed in *CHD8*^+/−^ human individuals. As such, the results offer insight into the neurodevelopmental pathology associated with mutations to *CHD8*, a genetic model that appears to be a bellwether for mutations affecting early transcriptional regulation and chromatin remodeling in autism. In-depth analysis of developmental neuroanatomy and social and communicative phenotypes as well as associated attentional and cognitive deficits in these *Chd8*^+/−^ mice will be necessary to link observed changes in brain gene expression and structure with relevant pathology in humans. This study of the impact of Chd8 haploinsufficiency *in vivo* in mice demonstrates the power of genomic and systems-level characterization of neurodevelopment in animal models towards resolving major questions about the genetic and neurodevelopmental origins of autism and intellectual disability.

## Experimental Procedures

See extended online methods for full description of experimental procedures.

### Generation of Chd8 mutant mice

We used Cas9-mediated mutagenesis of C56BL/6N oocytes to generate two mouse lines harboring frameshift deletions (5 bp and 14 bp) in mouse *Chd8* exon 5 (gRNA sequence: GAGGAGGAGGTCGATGTAAC). Guide RNA was synthesized and pooled with Cas9 mRNA^39^ and injected into mouse oocytes. We identified F0 pups carrying 5 bp and 14 bp deletions that overlap the target sequence. Heterozygotes were crossed to WT C57BL/6N mice to expand the lines and eliminate off-target mutations. We examined Chd8 protein and transcript levels via western blot (ab114126; Abcam) and qPCR at E14.5, P0, and adult forebrain and compared cortical length in whole mount P0 brains from *Chd8*^+/−^ mice and matched WT littermates. All mouse studies were approved by the Institutional Animal Care and Use Committees at the University of California Davis and the Lawrence Berkeley National Laboratory. Subject mice were housed in a temperature-controlled vivarium maintained on a 12 hour light-dark cycle. Efforts were made to minimize pain and distress and the number of animals used.

### Developmental neuroanatomy

Litters for neuroanatomy analysis were generated by breeding male *Chd8*^+/−^ mice with WT females. Brains were perfused before isolation, embedding, and sectioning. *P7 Nissl staining:* Nissl staining was performed following established protocols and morphological parameters were measured and compared using Student’s t-test. *E13.5 EdU labeling:* Timed-pregnant females were intraperitoneally injected with 50 mg/kg body weight EdU. After 1.5 hours, females were anesthetized and embryos were perfused, fixed, and sectioned. EdU detection was performed with the Click-it EdU Alexa 594 imaging kit protocol (Life Technologies) according to the manufacturer instructions. *P0 lamination:* Slides were incubated in blocking solution, rinsed in PBS-T, and incubated overnight at 4°C in primary antibody solution containing anti-Ctip2 (ab18465; Abcam) and anti-Tbr1 (ab31940; Abcam) antibodies. The slides were rinsed and incubated overnight at 4°C in fluorophore-conjugated secondary antibodies. Slides were counterstained for 2 hr in DAPI, rinsed, and mounted. All sections over the entire brain were surveyed for lamination defects and corresponding sections imaged. Within each genotype, all brains were selected randomly for histological processing without taking morphological criteria into account. All histology was done blind, by investigators that were unaware of group allocation. No data points were excluded. All antibodies used for this study were validated and their use widely reported.

### Genomics

Bulk forebrain was microdissected from *Chd8*^+/−^ and matched WT littermates at E12.5, E14.5, E17.5, P0, and from adults (>P56). Samples included males and females of each genotype at each stage. Total RNA was isolated using Ambion RNAqueous and assayed using an Agilent BioAnalyzer instrument. Stranded mRNA sequencing libraries were prepared using TruSeq Stranded mRNA kits and 6-12 samples per lane were pooled and sequenced on the Illumina HiSeq platform using a single end 50 bp (E145, E17.5, P0, adult) or paired end 100 bp (E12.5) strategy. FASTQ files were aligned to the mouse genome (mm9) and counts for mouse genes were calculated. For inclusion in testing, genes were required to have a minimum read count of at least 10 reads/million in more than two samples. Differential expression was performed with edgeR^40^ using generalized linear models including factor-encoded sex and developmental stage/sequencing batch as covariates. After normalization, iterative Weighted Gene Correlation Network Analysis (WGCNA^23^) was used to identify co-expressed gene modules. Fourteen discrete gene expression networks were identified (numbered by module gene count), and genes that were not classified into one of these modules were assigned to M.8.grey. Permutation testing was performed to test for overlap between DE genes and published gene sets. Gene Ontology Biological Process term enrichment and protein-protein interaction network analysis was performed using the TopGO Bioconductor package and STRING^26^. Differential expression of selected targets was verified by qPCR at P0. Primers reported in Table S3. For qPCR analysis, 9 wild-type and 7 *Chd8*^+/−^ forebrain samples were used. Samples were excluded if technical replicates failed. Paired t-test was performed on *Actb* normalized relative gene expression between WT and *Chd8*^+/−^ using ∆∆CT. To reduce noise, the highest and lowest values from both groups was discarded. Tbr1 protein level assayed via western blot (ab31940; Abcam) compared via Student’s t-test.

### MRI

After perfusion, brains from mice that underwent behavioral screening were scanned using a multi-channel 7.0 Tesla MRI scanner (Varian Inc., Palo Alto, CA). Diffusion Tensor Imaging (DTI) was done using a 3D diffusion weighted fast spin echo sequence to create fractional anisotropy, mean diffusivity, axial diffusivity, and radial diffusivity maps for brains used in this study. After registration, changes and intensity differences were examined for the volume or mean diffusion measure for 159 different structures encompassing cortical lobes, large white matter structures, ventricles, cerebellum, brain stem, and olfactory bulbs. Initially seven summary regions were examined, including the cerebral cortex, olfactory bulbs, cerebral white matter, cerebral gray matter, ventricles, brainstem, and cerebellum^41^. Multiple comparisons in this study were controlled for via False Discovery Rate.

### Behavioral testing

All procedures were approved by the University of California Davis Institutional Animal Care and Use Committee, and were conducted in accordance with the National Institutes of Health Guide for the Care and Use of Laboratory Animals. Efforts were made to minimize pain and distress and the number of animals used. No previous analyses were performed on animals used for behavioral testing. Subject mice were housed in a temperature-controlled vivarium maintained on a 12 hour light-dark cycle. We used mixed genotype home cages with 2-4 animals per cage and blinded experimenter and video scorer/processor to genotype during testing and analysis. All tests were conducted during the light cycle. Groups sizes indicated are based on past experience and power analyses. Effects of genotype and sex evaluated using Multi-factor ANOVA, as previously published. Significant ANOVAs followed with Tukey’s high significant difference test, or other appropriate post hoc tests including Bonferroni correction tests for multiple comparisons, to correct for false discovery and to identify specific differences between groups. Behavioral analysis passed distribution normality tests, was collected using continuous variables and thus analyzed via parametric analysis, in all assays. For all behavioral analyses, variances were similar between groups and no data points were excluded. One animal died during behavioral testing. This happened during Three Chamber Social approach, in the middle of the behavioral battery. *Chd8*^+/−^ male and female mice and WT littermates, ages 2-5 months, were evaluated in a standard battery of neurobehavioral assays relevant to diagnostic symptoms of autism^27^. In total, 8 male and 10 female *Chd8*^+/−^ mice and 11 male and 10 female matched WT littermates were tested in the following sequence: open field, general health, self-grooming, marble burying, 3-chambered social approach, and male-female social interactions. Testing was performed at the UC Davis MIND Institute Intellectual and Developmental Disabilities Research Center Mouse Behavior Core. Statistical testing was performed using established assay-specific methods, including Students t-test for single parameter comparisons between genotypes, and One-Way or Two-Way Repeated Measures Analysis of Variance for comparisons across time points and/or between sexes.

### Data availability

All relevant data will be available from authors. DOIs for all published gene sets used in enrichment analysis: Sanders et al. 2015 - 10.1016/j.neuron.2015.09.016; Parikshak et al. 2013 - 10.1016/j.cell.2013.10.031; Cotney et al. 2015 - 10.1038/ncomms7404; Willsey et al. 2015 - 10.1016/j.cell.2013.10.020; Sugathan et al. 2014 - 10.1073/pnas.1405266111; Darnell et al. 2011 - 10.1016/j.cell.2011.06.013; Hormozdiari et al. 2014 - 10.1101/gr.178855.114; Voineagu et al. 2011 - 10.1038/nature10110.

### Code availability

All custom scripts used for data processing and analysis will be available from authors. A custom sample processing pipeline was used to align raw sequencing samples to mouse genome mm9 using RNA-seq aligner STAR (version 2.4.2a), features assigned via subreads featureCounts (version 1.5.0) to UCSC mm9 genes.gtf, and quality check performed on individual samples using RSeQC (version 2.6.3). Differential expression analysis was done with a custom pipeline in R Studio using functions from edgeR (version 3.10.5) and limma (version 3.24.15). Permutation testing was performed with a custom R script. Iterative co-expression network analysis was performed with a custom pipeline following the standard WGCNA (version 3.2.3) workflow and functions. Gene Ontology analysis was performed with a custom wrapper using standard the TopGO (version 2.20.0) program. See extended methods for description and parameters.

### Author Contributions

ALG, JE, JPL, JNC, JLS, KSZ, and ASN designed the experiments. Generation of mouse model: ASN, DD, AV, LAP, BM, IPF, VA; Mouse behavior: NAC, MCP, MDS, JNC, JLS; Mouse MRI: JE, JPL; Genomics: ALG, LS-F, IZ, BM, ASN; Neuroanatomy: ALG, TWS, IZ, KSZ. ALG, LS-F, JE, KSZ, JC, JLS, and ASN drafted the manuscript. All authors contributed to manuscript revisions.

## Acknowledgements

Sequencing was performed at the UC Berkeley and UC Davis DNA cores. This work was supported by institutional funds from the UC Davis Center for Neuroscience and by the UC Davis MIND Institute Intellectual and Developmental Disabilities Research Center (U54 HD079125). L.S.-F. was supported by the UC Davis Floyd and Mary Schwall Fellowship in Medical Research. A.V., L.A.P, and D.E.D. were supported by National Institutes of Health grants R24HL123879, U01DE024427, R01HG003988, U54HG006997, UM1HL098166. Research conducted at the E.O. Lawrence Berkeley National Laboratory was performed under Department of Energy Contract DE-AC02-05CH11231, University of California.

